# Impactful feeding ecology of a predatory guild of oceanic jellies

**DOI:** 10.1101/2024.08.19.608650

**Authors:** Thomas Irvine, John H. Costello, Brad J. Gemmell, Kelly R. Sutherland, Sean P. Colin

## Abstract

Planktonic organisms are integral members of open ocean ecosystems and are critical drivers of material cycles in the world’s oceans. Ctenophores may be numerically dominant predators in these oceanic ecosystems but have been ignored due to the difficulty in both sampling and handling their extremely delicate, gelatinous bodies. To better understand their trophic impact, we combined SCUBA with novel imaging techniques to non-invasively document prey ingestion patterns of four widespread oceanic ctenophore species. We found that these ctenophores ingested 32 prey per hour and the most voracious species ingested nearly 50 prey per hour. Further, the size and number of prey ingested increased with ctenophore size. At these rates, lobate and cestid ctenophores consume prey at similar rates to their highly impactful coastal relative, *Mnemiopsis leidyi* and are likely the most impactful planktonic predator in the open oceans. Further, we showed that although major dietary components overlapped, different oceanic ctenophore species appear to specialize on different members of the plankton. Since these oceanic ctenophore species frequently co-occur, they comprise a powerful guild of influential planktonic predators with synergistic impacts. These results indicate that epipelagic ctenophores have much greater trophic effects on material cycles over broad areas of the open ocean than previously considered. Models of oceanic carbon cycling will benefit by more fully incorporating the impacts of oceanic ctenophores on their planktonic prey.

## Introduction

The emergence of optical plankton sampling techniques has revealed that gelatinous zooplankton predators dominate the biomass of global oceanic ecosystems ^1^. Conventional net sampling techniques, which have been used for centuries, inadequately sample gelatinous species and, in particular, they severely under-sample ctenophores (commonly called comb jellies; ^2–5^). Updated estimates have found that gelatinous zooplankton predators make up 30% of total biovolume, and 10% of the carbon within oceanic plankton from the tropics to the poles ^1^. This recognition of the predominance of gelatinous zooplankton challenges our understanding of oceanic food-web dynamics because gelatinous zooplankton have been viewed as only a minor component of oceanic systems and only minor players in oceanic biogeochemical cycles which are critical in regulating atmospheric carbon dioxide ^1,6–9^.

A consequence of historical under-sampling of oceanic ctenophores has been that they have generally been left off the list of organisms considered within oceanic nutrient flux models. In tandem with improved abundance estimates of ctenophores, non-invasive *in situ* techniques are elucidating the feeding mechanics of oceanic ctenophores ^10–13^. These studies indicate that oceanic lobate ctenophores use prey encounter and capture mechanisms similar to the coastal lobate ctenophore *Mnemiopsis leidyi. M. leidyi* has proven a highly invasive species throughout Europe and is more accessible to shore-based laboratories. For these reasons, it has been studied much more extensively than any other ctenophore species. The resulting body of research has unambiguously demonstrated *M. leidyi* predation can limit zooplankton populations and, consequently, produce cascading effects on trophic groups as distant as phytoplankton and bacteria ^14–17^.

While some ctenophores, such as *M. leidyi*, have been shown to be voracious predators, feeding mechanisms and diets vary greatly among genera. Lobate ctenophores use fused cilia, or ctenes, to swim and generate feeding currents to initiate encounters with prey. The most studied lobate ctenophore, *M. leidyi*, actively scans their feeding current to detect prey prior to physical contact ^18^. Cestid ctenophores, such as the Venus Girdle (i.e.; *Cestum veneris*), also swim to encounter prey, however, they lack the auricles and lobes of the oceanic lobate species. Instead, cestid ctenophores rely on prey colliding with tentillae that blanket their wing-like body as they swim through the water ^19^. Finally, cydippid ctenophores, like those of the genus *Pleurobrachia*, are ambush predators and have a passive feeding strategy that relies on prey swimming into their extended tentillae ^10^. The active feeding mechanism of lobates, and likely cestids, results in much higher encounter and feeding rates than cydippid ctenophores ^20,21^.

Multiple species of lobate and cestid ctenophores commonly co-occur in the oceanic surface waters around the globe. Recent studies have shown that this group of predators process as much fluid and encounter and capture prey at similar rates to the coastal lobate, *M. leidyi* ^12,13,22^). These studies suggest that oceanic ctenophores may exert a trophic impact comparable to the coastal *M. leidyi*, however, very few studies have quantified feeding by open ocean ctenophores.

In this study, we use novel, non-destructive *in situ* imaging methods to quantify the gut contents of common, cooccurring lobate and cestid oceanic ctenophores. These methods provided a rare glimpse into the gut contents of these extremely delicate pelagic predators and enabled us to quantify the feeding selection of about 300 pelagic ctenophores across four common taxa: the lobates *Bolinopsis vitrea., Eurhamphaea vexilligera,* and *Ocyropsis* spp., and the cestid *Cestum veneris* (Fig. 1A-D).

**Figure 1-.**
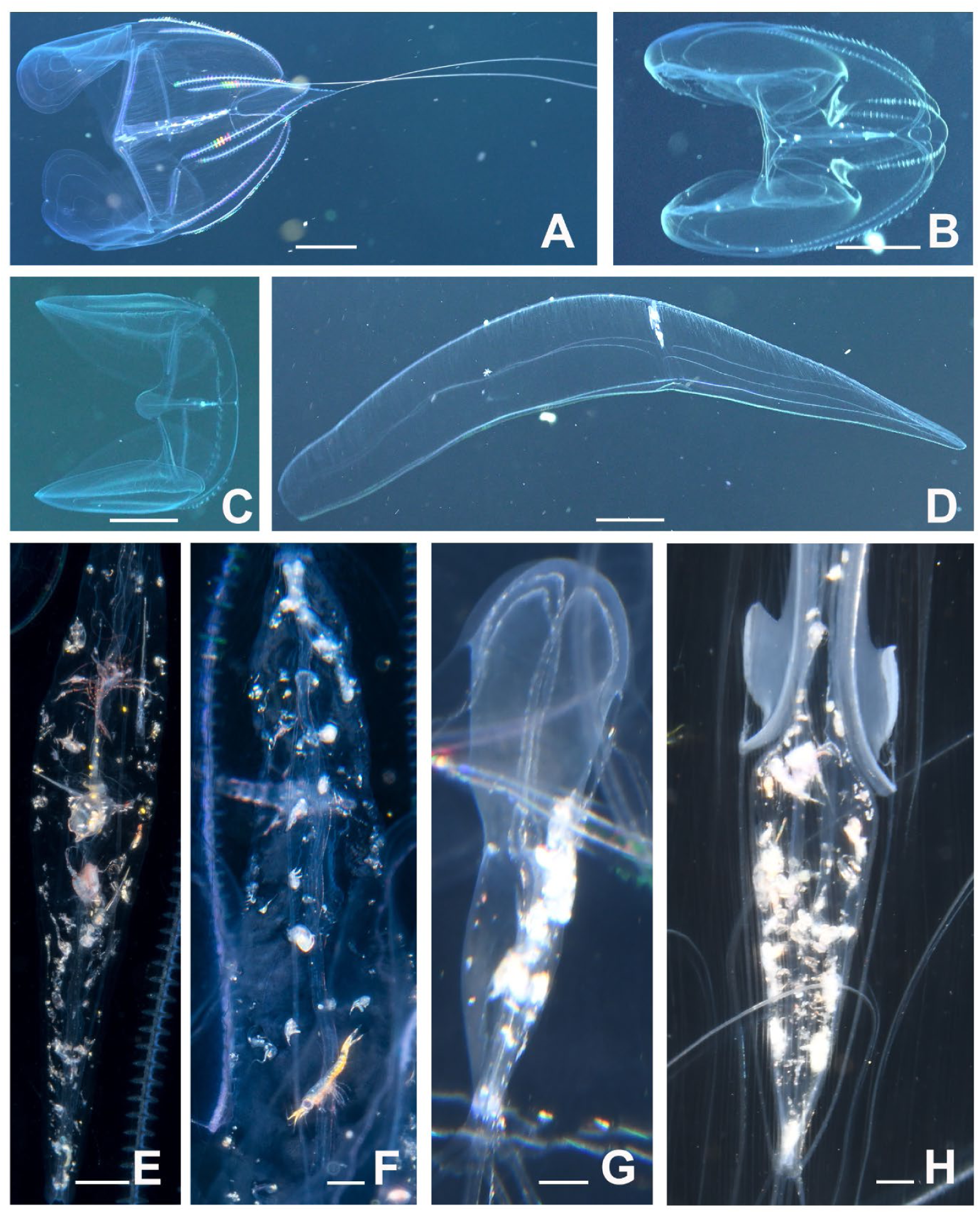
Images of common, cooccurring oceanic lobate and cestid ctenophores (A-D) and their guts with different prey items visible (E-H). A, E) *Eurhamphaea vexillegera*; B, F) *Bolinopsis vitrea*; C, G) *Ocyropsis crystallina*; D, H) *Cestum veneris*.

## Methods

### Image collection

*In situ* ctenophore images and videos were taken during drift dives at West Palm Beach, FL (26⁰ 43’ 93” N, 79⁰ 59’ 15” W) and Kona, HI (19° 40’ 10.5” N 156° 02’ 46.4” W) from 2021 to 2023. A total of 299 ctenophores were analyzed, including 47 *Bolinopsis vitrea*, 64 *Cestum veneris*, 93 *Eurhamphaea vexilligera*, and 95 *Ocyropsis* spp. The gut contents of these oceanic ctenophores were recorded from either photographs or high-resolution video of the ctenophore drifting in their natural setting. When imaged properly, the contents of their guts are visible through the transparent ctenophore tissues (Fig. 1E-H; Rapoza et al., 2005). Video images were collected using high-resolution 4K video cameras (Sony AX100 and Z-cam E2-M4) with brightfield collimated-light optical systems ^13,24^. Still photographs were taken using DSLR cameras with strobe flash setups. Photographs were taken by both our research group and by citizen scientists who are underwater photographers that regularly participate in blackwater drift dives in West Palm Beach, FL (see Acknowledgements for specific individuals). Stills or video frames used for gut content analysis met the following requirements: 1) the entirety of the gut was in frame and, 2) contents of the gut were in focus and unobscured.

### Image analysis and gut content analysis

The following measurements were taken for all ctenophore images using Fiji ImageJ (NIH): 1) total prey count in the gut, 2) gut area, 3) individual prey item area, length and width and 4) prey type for identifiable items. Images were scaled by placing a ruler in the field of view during image collection. Gut fullness was calculated as the total area of the prey divided by the total area of the gut ^12^. Prey length was taken as the distance from the anterior to the posterior end of the prey when the anatomical features were discernible, otherwise the length was the longest distance across the prey item. Width was taken as the perpendicular measurement halfway along the length measurement (Fig. 1).

Ingestion rate (*I*) was estimated as the total number of prey in the gut divided by the digestion time (*t*). We used 44 minutes as the digestion time, and was based on measurements for *O. crystallina* fed different amounts of copepod prey at 25°C ^12^. The Potter et al. (2023) temperature was several degrees cooler than the water conditions of our studies and, therefore, represents a conservative estimate of digestion time. Carbon consumption rates were estimated using the equation:

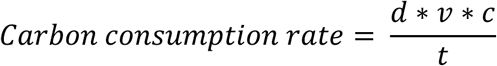

Where total prey biovolume (*v*) was calculated as the sum of the prey volume estimates where individual prey volumes were estimated assuming prey were ellipsoidal with the major and minor axis being the prey length and width measurements, respectively. The density (*d*) of individual prey items was assumed to be 1.0273 mg mm^−3^ ^12^ and the carbon per unit wet-weight (*c*) was assumed to be 0.0684 mgC mgWW^−1^ of prey (Madin et al., 2001, Potter et al. 2023). For lobate ctenophores where the entire predator was in-frame, ctenophore lengths were measured from the point between the oral lobes to the posterior end of the ctenophore. Length of *C. veneris* was recorded as the length from the mouth to the statocyst. In the case of a few of the outsourced images, ctenophore measurements were recorded while *in situ*.

Prey diversity was assessed by documenting all the identifiable prey taxa in the guts (*B. vitrea* (n = 20), *C. veneris* (n = 43), *E. vexilligera* (n = 45), and *O. crystallina* (n = 33)). The Shannon Diversity Index (H’) of the gut contents was calculated as:

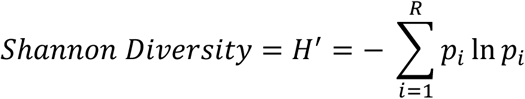

Where *p_i_* is the proportion of individual prey items belonging to the *i*th species and R is the number of species/species groups.

### Statistical Analysis

Differences among species were compared using One-way ANOVA tests to determine whether there were significant differences in prey count, gut fullness, prey length, and carbon consumption estimates. Dunn’s Method was used for Post-hoc analysis of significant ANOVA tests. All data were tested for homoscedasticity and normality. If the data did not conform to the assumptions of an ANOVA, then a Kruskal-Wallis Ranks test was used. Linear regressions were used to determine if there were any significant correlations between the length of the ctenophore and prey count, prey length, and estimated carbon consumption rates.

## Results

Using non-invasive SCUBA-based methods to quantify gut contents of planktonic organisms in their natural environment is difficult and time consuming. In order to collect sufficient observations to obtain reliable quantitative gut content data, we collected data over many days and multiple years. Therefore, our observations represent averages over highly variable environmental, prey and predator conditions. Despite the data representing relatively longer-term averages, we observed significant differences in the amount and sizes of the prey found in the guts of the different lobate and cestid ctenophores (Fig. 2A and B; Kruskal-Wallis Ranks test, p < 0.01). *Cestum veneris* and *Eurhamphaea vexilligera* consumed the most prey, having on average 29.8 and 31.6 prey gut^−1^, respectively (Fig. 2A; Dunn’s Method Post-hoc analysis, P < 0.001). With an assumed conservative digestive time of 44 minutes ^12^, this translates to estimated ingestion rates of 45.2 and 47.9 prey hr^−1^ respectively. *Bolinopsis* and *Ocyropsis* spp. consumed less prey with *Ocyropsis* spp. consuming the least amount of prey (Dunn’s Method Post-hoc analysis, P < 0.005). In addition to consuming the most prey, *C. veneris* also ingested the largest prey (Fig. 2B; Dunn’s Method Post-hoc analysis, P < 0.005). The more abundant and larger prey in the guts of *C. veneris* translated to it having an average carbon consumption rate of 1.6 mgC hr^−1^, more than double the rate of the other ctenophores (Fig. 2D, Dunn’s Method Post-hoc analysis, P < 0.01).

**Figure 2-.**
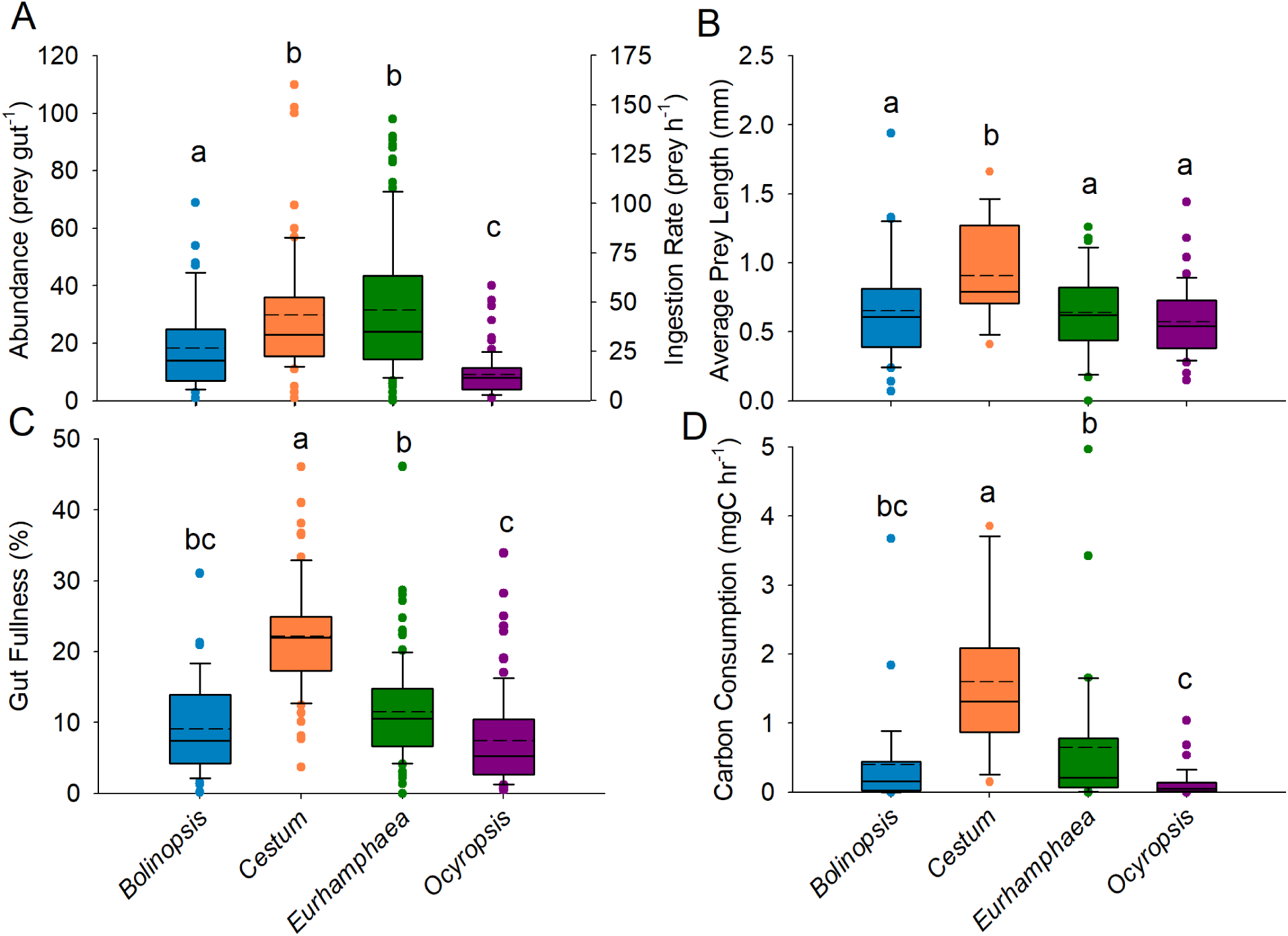
Comparison of prey size and abundance in the guts of co-occurring ctenophores. A) Abundances, ingestion rates and B) lengths of prey in the guts of *Bolinopsis vitrea* (n=47), *Cestum veneris* (n=64)*, Eurhamphaea vexilligera* (n=93) and *Ocyropsis crystallina* (n=93). C) Gut fullness and D) carbon consumption rates of the four ctenophore species. Boxplots indicate interquartile ranges, mean is indicated by the dashed line and median by the solid central line. Black circles show outlying data points. Lowercase letters denote significantly different values (Dunn’s Method Post-hoc analysis, P < 0.05).

Gut fullness was determined by comparing the area of the gut containing prey to the total 2-dimensional gut area (ignoring any 3-dimensionality of the guts). Based on this comparison, the guts of *C. veneris* were fuller than the other ctenophores (Fig. 2D, Dunn’s Method Post-hoc analysis, P < 0.01), however, the mean gut fullness of all the species was less than 25% and no guts observed were greater than 50% full (Fig. 2C). This suggests that these species have the gut capacity to utilize dense patches of prey they may encounter in their environment.

The number and size of prey consumed and the estimated carbon consumed by *C. veneris* and *E. vexilligera* significantly increased with ctenophore size (Fig. 3A and B; Regression Analysis, P < 0.05). Notably, large *E. vexilligera* (> 7 cm) had more than 5 times the number of prey in their guts than average sized *E. vexilligera* (3.7 cm). Since prey were also larger for larger ctenophores, large *E. vexilligera* could consume greater than 6 times the carbon hr^−1^ than average sized *E. vexilligera* and large *C. veneris* could consume more than 8 times the carbon hr^−1^ of averaged sized conspecifics. Interestingly, the abundance of prey in the guts of the oceanic ctenophores in this study were equal to and even greater than the abundance of prey in the guts of *Mnemiopsis leidyi* from both the Narragansett Bay, RI, USA (Costello & Sullivan, unpublished data) and Gullmar fjord, Sweden ^20^.

**Figure 3-.**
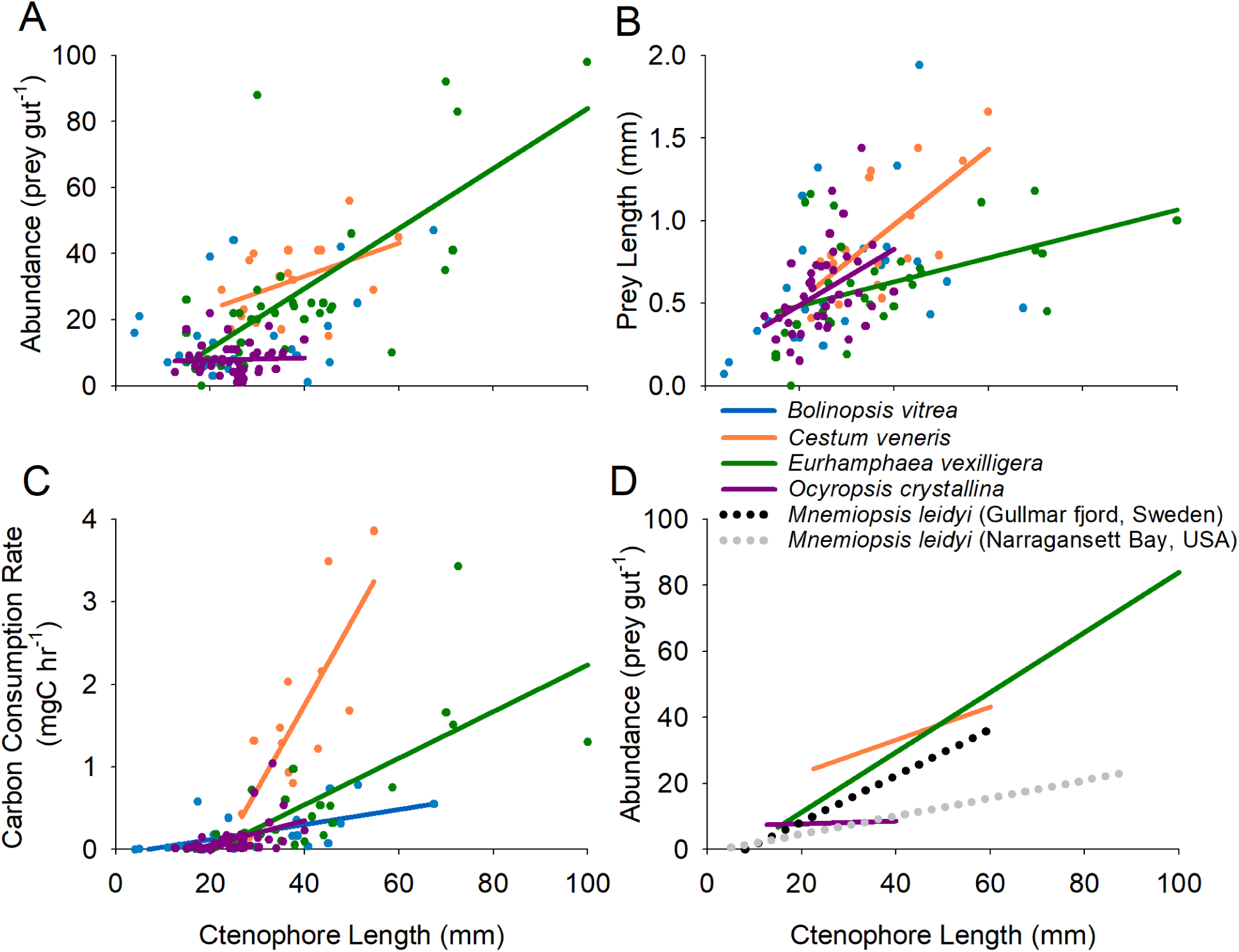
Effects of ctenophore size on the gut contents of the cooccurring ctenophores. A) Abundance and B) size of prey in guts of different sized *Bolinopsis* vitrea (n=47), *Cestum veneris* (n=64)*, Eurhamphaea vexilligera* (n=93) and *Ocyropsis crystallina* (n=95). C) Carbon consumption rates of different sized ctenophores. The amount and size of prey increase with ctenophore size for all species except *B. vitrea*. D) Trendlines of prey abundance (same as A)) in guts of oceanic ctenophores compared to coastal lobate ctenophore *Mnemiopsis leidyi* (dotted lines) from two locations show that oceanic ctenophores ingest as many prey as their nearshore relative, *M. leidyi*. Solid trend lines were included for genera where the trends were significant (Regression analysis, p<0.05).

*E. vexilligera* consumed a more diverse assemblage of prey than the other ctenophore taxa (Fig. 4A) with > 20 different prey types observed in the guts (Table S1). While copepods dominated the guts of all the ctenophores, *E. vexilligera* commonly had pteropods, tintinnids and even fish in their guts (Table S1, Fig. 4C). In contrast, *Ocyropsis* spp. consumed almost exclusively copepods and as a result had the least diverse prey assemblage in their guts. However, the occasional pteropod, tintinnid and other invertebrate were seen in the guts of all species (Table S1, Fig. 4C). In addition to prey types, the size distribution of consumed prey differed among species. The mean size of prey consumed by all species was between 0.5-1 mm (Fig. 2D and Fig 4D). The sizes of the prey consumed by *Ocyropsis* spp. were least variable among the ctenophores while there were multiple size peaks in the prey distribution for *E. vexilligera* and *C. veneris* (Fig. 4D). As a community, the ctenophores consumed almost 25 different types of taxa (Fig. 4B) and with a broader range of sizes than each individual ctenophore species.

**Figure 4-.**
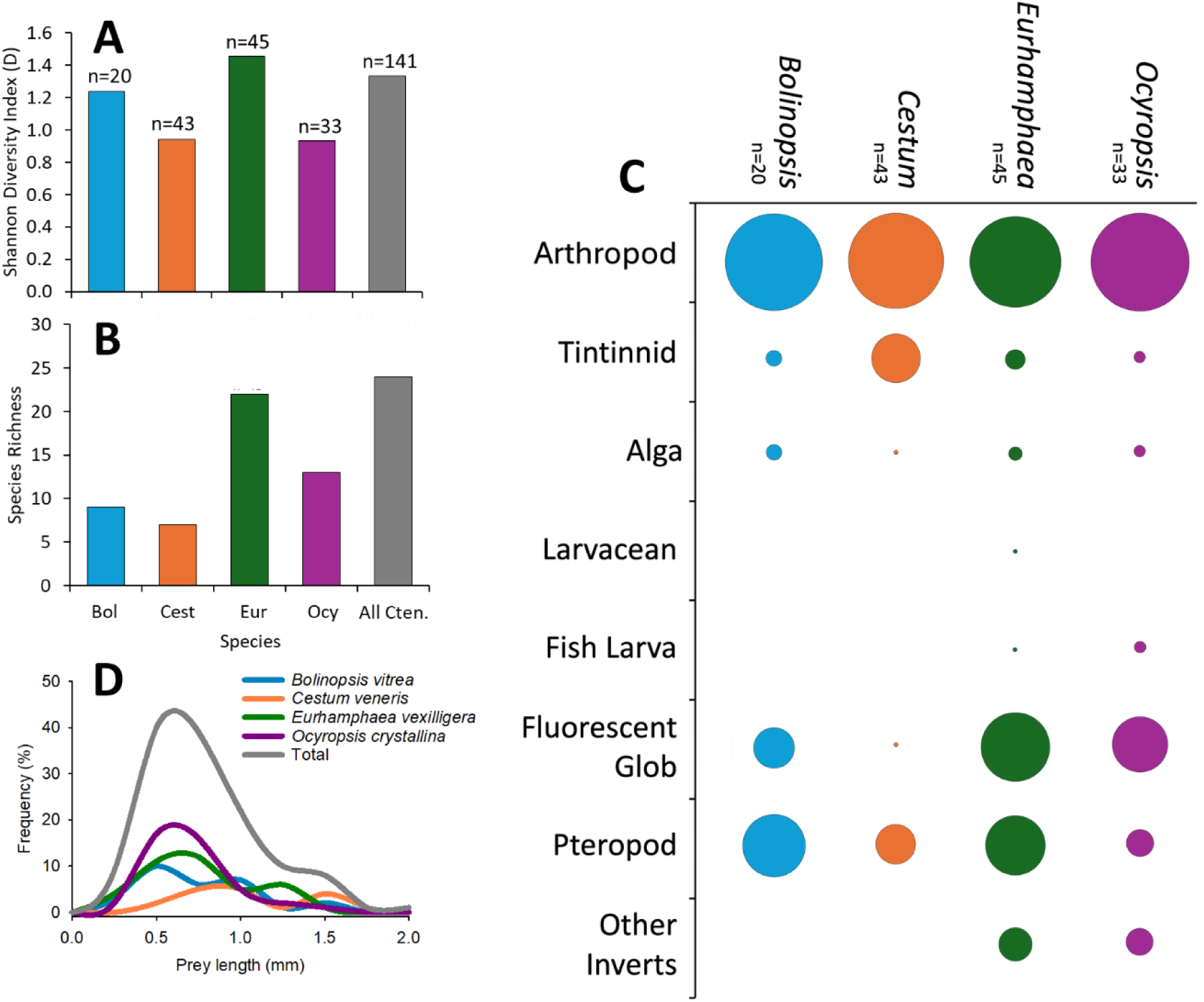
Comparison of diet diversity among oceanic ctenophores. A) Shannon Diversity Index and B) species richness both show differences in how diverse the diets were among *Bolinopsis vitrea, Cestum veneris, Eurhamphaea vexilligera* and *Ocyropsis crystallina*. C) Bubble diagram of log proportionality of the different prey species identified in the guts show that while copepods dominated all the guts, different taxa were more abundant in the guts of the different ctenophores. “Arthropod” (panel C) was mostly copepods but a small number of decapods and ostracods were also observed). In addition to prey taxa, D) average prey sizes per gut varied among species as is illustrated in the relative frequency distribution of prey size.

## Discussion

More comprehensive sampling using new optical techniques are revealing that the oceanic pelagic ecosystems are jelly-dominated habitats ^1,25^. A diverse range of taxa comprise gelatinous zooplankton communities and include cnidarians such as siphonophores and medusae, tunicates such as salps and larvaceans, mollusks such as pteropods and heteropods, chaetognaths, and ctenophores. Among this broad gelatinous group, we found that lobate and cestid ctenophores have notably high consumption rates with some species consuming an average of about 50 prey per hour. The number of prey in the guts of oceanic ctenophores studied were similar to, and even greater than, the number of prey found in the guts of *Mnemiopsis leidyi* (Fig. 3D), a well-documented, dominant zooplankton predator in coastal ecosystems. These high prey counts were observed despite oceanic waters surrounding these ctenophores being considerably less productive than the coastal waters where *M. leidyi* is found. In addition, the abundance of prey in the guts of the oceanic ctenophores were several orders of magnitude greater than have been quantified for other oceanic gelatinous predators (Table S2).

The high predatory impact of lobate and cestid ctenophores is enabled by their feeding mechanisms that rely upon feeding-currents to stealthily process large volumes of fluid ^13,26^ and retain prey with high efficiency ^18,22,27^. In order to feed, the lobate and cestid ctenophores use their ctene rows to swim slowly through the water column and to transport fluid past their sensory and capture surfaces ^10,11,13,26,28^. Despite lacking eyes, these ctenophores are capable of sensing and reacting to the presence of prey items and capturing most prey they encounter ^12,27^. An advantage of feeding without vision is the ability to feed throughout the night as well as during daylight hours. This strategy has enabled *M. leidyi* to cause trophic cascades ^29–31^ and out compete fish larvae prey in coastal ecosystems ^32,33^. In oceanic systems, this feeding strategy enables oceanic lobate and cestid ctenophores to encounter > 2 prey per minute and ingest >40% of these prey ^22^. These estimates of ingestion rates based on encounter rates and retention efficiencies ^22^ support our estimates of ingestion rates based on gut contents (48 versus 47 prey h^−1^, respectively, when considering *Eurhamphaea vexilligera* and *Cestum veneris*). At these rates, oceanic lobate and cestid ctenophores consume several orders of magnitude more prey than all other ambush-feeding gelatinous predators (Table S2). These stealthy, feeding-current strategies have enabled lobate and cestid ctenophores to be probably the most impactful gelatinous predators in the oceans.

Since lobate and cestid ctenophores encounter and sense prey using similar mechanisms, it is not surprising to find they have overlapping diets and all consume the ubiquitous copepods found in epipelagic surface waters. However, we also observed distinct differences in the types and size of prey ingested. For example, *E. vexilligera* and *C. veneris* ingested larger prey than the other species and *E. vexilligera* also ingested more types of prey. *Ocyropsis* spp. appeared to specialize on copepods while *Bolinopsis vitrea* favored smaller prey. A consequence of having overlapping but different diets is that, as a community of predators, lobate and cestid ctenophores will have a cumulative impact on many species (especially copepods) and have a broader impact on the ecosystem. Unlike *M. leidyi*, which is typically the lone lobate ctenophore species in the coastal ecosystems, these oceanic species co-occur in oceanic ecosystems worldwide ^34–37^ and, therefore, represent a widely distributed guild of predators. An accurate assessment of their impact requires consideration of the full ctenophore guild’s synergistic effects on planktonic communities.

Perhaps surprisingly given their demonstrably high feeding rates, the predatory impact of this guild is unknown. Conventional net sampling methods greatly under sample lobate and cestid ctenophores ^3,5,38^ and there have not been enough studies published to date that have used optical samplers to quantify ctenophore abundances and changes in abundances over time and space. Potter et al. (2022) estimated, based on gut content analysis, that at relatively high densities *Ocyropsis* spp. alone was capable of removing > 40% of the copepod standing stock per day, while at low densities they removed only < 1%. Our data of *Ocyropsis* spp. was very similar to the ingestion rates estimated by Potter et al. (2022) but *Ocyropsis* spp. had the lowest ingestion rates among the species we examined. Therefore, the cumulative impact of the entire ctenophore predatory guild will be greater than estimated for *Ocyropsis* spp. Their collective tropic impact ultimately depends on ctenophore abundances and, due to the challenges of oceanic ctenophore’s body designs, these data are sparse and/or underestimate abundance. However, in order to arrive at a first order approximation of the ctenophore abundances necessary to generate a significant impact on the zooplankton community, we can use our measured community-wide ingestion rates – i.e.; mean ingestion rates among all species – in conjunction with average epipelagic zooplankton prey densities from around the world to estimate the half-life of zooplankton populations. The half-life of zooplankton populations (*t_1/2_*) is the time it takes predators to reduce the concentration of zooplankton prey (C, prey m^−3^) by 50% ^16,39^. It is calculated as:

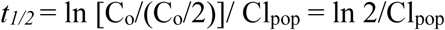

Where Cl_pop_ (m^3^ d^−1^) is the clearance rate of the ctenophore population and Co/(Co/2) = 2 when the initial prey concentration (C_o_) is reduced by 50%. The clearance rate of the ctenophore population is:

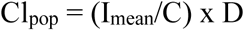

Where I_mean_ is the average ingestion rate (prey ctenophore^−1^ day^−1^) among all the ctenophore species, C is the prey concentrations (prey m^−3^) and D is the concentration of ctenophores (ctenophores m^−3^). The two unknowns in estimating *t_1/2_* are the abundance of ctenophores (D) and prey (C). However, using this concept of zooplankton half-life we can then estimate the ctenophore abundances necessary to reduce C by ½ every day – i.e.; *t_1/2_* = 1 day – for different zooplankton abundances (Fig. 6). We use *t_1/2_* = 1d because it corresponds to peak predatory impact rates recorded for *M. leidyi* in coastal ecosystems ^40–42^ and represents extreme predation pressure that could quickly decimate zooplankton populations. The drop lines in Fig. 6 indicate different abundances of epipelagic zooplankton (Table S3) from global averages (solid drop lines) and from different tropical/sub-tropical locations from around the world (dashed drop lines). Based on our measured ingestion rates it would take less than 0.04 ctenophores m^−3^ (or 40 ctenophores1000 m^−3^) to ingest half the zooplankton stock on a daily basis based upon global averages of epipelagic zooplankton abundances ^43^. Further, estimates of epipelagic zooplankton from around the world required considerably less than 1 ctenophore m^−3^ to impact prey with a *t_1/2_* = 1 day.

**Figure 6.**
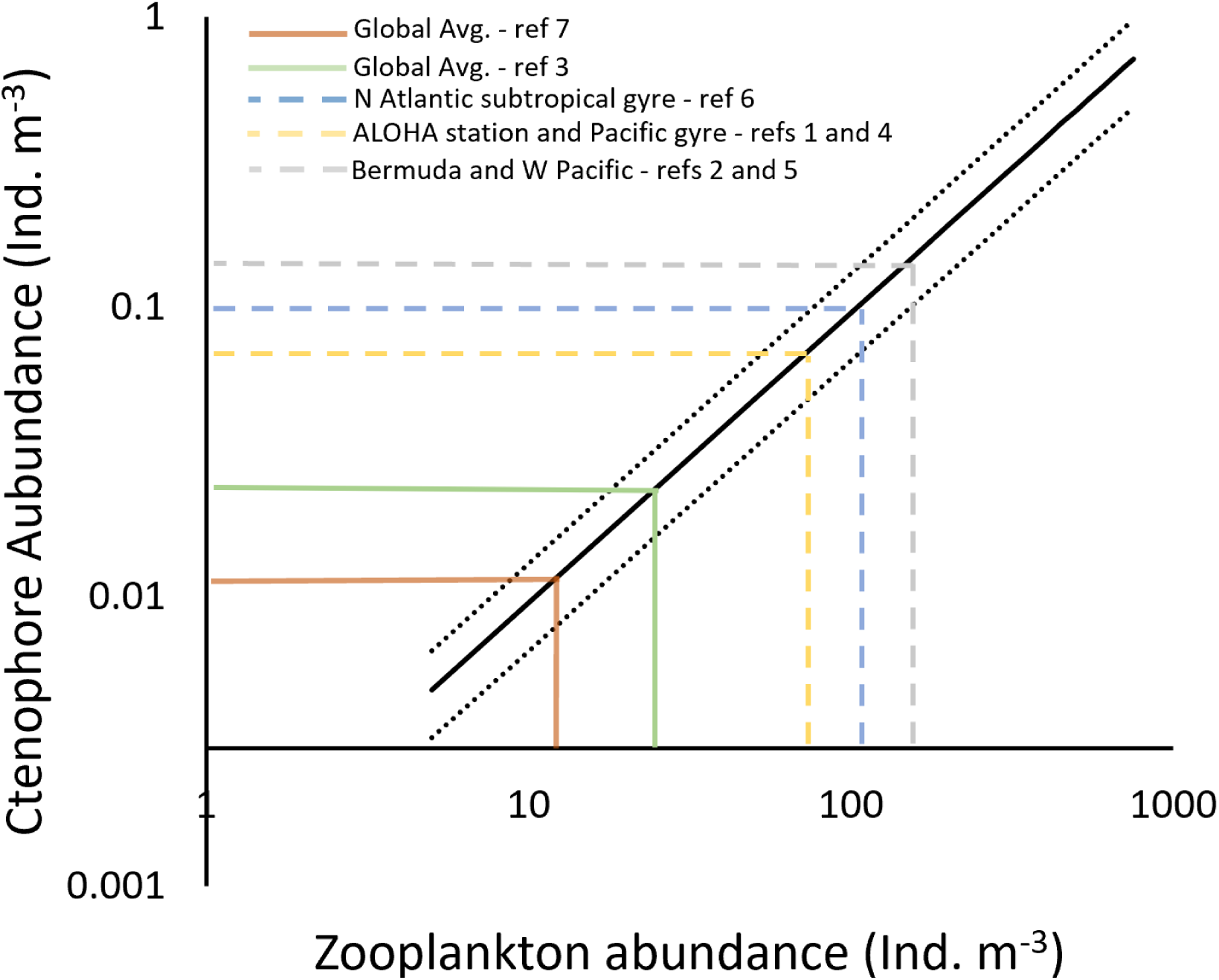
Estimated ctenophore abundances needed to ingest half of the zooplankton population per day (t_1/2_ = 1 day). Black solid line is based on ctenophore ingestion rates calculated using a digestion time of 44 minutes (Potter et al. 2023). Dotted black lines are high and low ingestion rates based on 60 and 30 min digestion times, respectively. Colored drop lines indicate ctenophore abundances needed for t_1/2_ = 1 for different global estimates (solid) and measured mesozooplankton abundances (dashed). Table S3 has more information on references.

Unfortunately, without reliable estimates of ctenophore densities it is not possible to evaluate how often ctenophore reach abundances that enable them to seriously impact zooplankton standing stocks. However, a seminal ctenophore study from Harbison et al. (1978) that involved diving in > 250 oceanic locations around the world found ctenophore abundances to be extremely variable ranging from zero ctenophores to much greater than 1 m^−3^. In fact, 5% of their locations had ctenophore concentrations greater than 1 m^−3^. At this density the ctenophore predatory guild is capable of impacting zooplankton at the global average (40 m^−3^) with a *t_1/2_* = 0.9 hours.

## Conclusion

The oceanic lobate and cestid ctenophores examined here co-occur globally in subtropical and -tropical epipelagic waters around the world ^2^ and make up a predatory guild that are arguably the most impactful zooplankton predators in epipelagic ecosystems. While they are primarily crustacean predators, they have broad and overlapping diets that regularly include tintinnids, pelagic mollusks, radiolarians, larvaceans and fish larvae. Ctenophores do not need to be highly abundant to greatly impact zooplankton standing stocks because the high prey capture efficiencies of their feeding-current mechanics allow a low concentration of oceanic ctenophores to have a large effect. Reasonable estimates of their feeding rates and plankton distributions indicates that oceanic ctenophores may regularly occur at densities that decimate zooplankton populations in hours. However, in order to evaluate the trophic consequence of their high predatory rates, it is clear that substantially more work needs to be invested in quantifying ctenophore abundances in oceanic systems. Despite scant abundance data, the current evidence indicates that oceanic lobate and cestid ctenophores consume enough prey to be important players in global oceanic ecosystems and global carbon cycling. Focused work on ctenophore distributions will allow accurate assessment of the actual magnitude of their impact.

