## Supplementary Tables for "Impactful feeding ecology of a predatory guild of oceanic jellies"

**Table S1**. Prey species observed in the guts of four pelagic ctenophores. Gray highlighted rows indicate species that were present in the diets of all four ctenophore species. Sample sizes: *Bolinopsis*=20, *Eurhamphaea*=45, *Ocyropsis*=33, *Cestum* = 43

| | Ctenophore taxa | | | |
|---|---|---|---|---|
| Prey species | *Bolinopsis vitrea* | *Cestum veneris* | *Ocyropsis crystallina* | *Eurhamphaea vexilligera* |
| Amphipod | X | X | X | X |
| Chaetognath | | | | X |
| Ciliate | | X | | X |
| Cladoceran | | X | | X |
| Copepod | X | X | X | X |
| Crab zoea | | | | X |
| Cyddipid larva | | | X | |
| Decapod larva | X | X | X | X |
| Diatom | | | | X |
| Dinoflagellate | X | | | |
| Fish larva | | X | X | X |
| Florescent glob | X | X | X | X |
| Gastropod larva | | | | X |
| Invertebrate egg | | | X | X |
| Larvacean | X | X | | X |
| Medusoid | | | | X |
| Mollusc larva | | | | X |
| Nematode | | | | X |
| Ostracod | X | X | X | X |
| Polychaete larva | | | | X |
| Pteropod | X | X | X | X |
| Radiolarian | | X | | X |
| Salp | | | | X |
| Tintinnid | X | X | X | X |

**Table S2.** Ingestion rates of epipelagic gelatinous predators based on gut content analysis of field collected specimens. Some studies did not measure ingestion rates but recorded percent of predators with prey in their guts.

| Taxa | Species | Ingestion Rate (prey/hr) | % pred. with prey in guts | Source |
|---|---|---|---|---|
| siphonophore | *Muggiaea* | 0.330 | 65 | Purcell 1982 |
| chaetognath | *Sagitta elegans* | 0.229 | 50 | Saito and Kiorboe 201 |
| chaetognath | *S. enflata* | 0.528 | 0.15 | Kehayias et al 2005 |
| chaetognath | *S. minima* | 0.028 | 0.5 | Kehayias et al 2006 |
| chaetognath | *S. Setosa* | 0.010 | 0.33 | Kehayias et al 2007 |
| chaetognath | *S. serratodentata* | 0.038 | 0.25 | Kehayias et al 2008 |
| chaetognath | All species | 0.603 | | Kehayias et al 2009 |
| heteropod | *Carinaria cristata* | | 0.4 | Seapy 1980 |
| hydromedusae | *Solmissus* | | 0.0205 | Raskoff 2002 |

**Table S3.** Mesozooplankton prey abundances at different tropical/subtropical locations around the world. The colors and reference numbers correspond to Fig. 5.

| Latitude | Longitude | Location Name | Taxa | Prey Density (ind/m$^3$) | % Copepods | Ctenophore density (ind/m3) | Reference number | Reference | Notes |
|---|---|---|---|---|---|---|---|---|---|
| 22 45 N | 158W | ALOHA station - subtropical gyre | mesozooplankton | 73.5 | 72.3 | 0.0706 | 1 | Steinberg et al 2008 | mean of high and low values of 67 and 81 |
| 35N | 155E | Western Pacific | mesozooplankton | 70 | 80 | 0.0672 | 2 | Yamaguchi et al 2017 | 10 year avg |
| 35 45 | 155E | Western Pacific | mesozooplankton | 150 | 75 | 0.1440 | 2 | Yamaguchi et al 2017 | 10 year avg |
| | | Global | mesozooplankton | 18 | | 0.0173 | 3 | Moriarty et al 2013 | |
| 15N-15S | | tropical | mesozooplankton | 12 | | 0.0115 | 4 | Moriarty et al 2013 | to depth of 350m |
| 32 45N | 64 30 W | Bermuda | mesozooplankton | 160 | 78.4 | 0.1536 | 5 | Stefanoudis et al 2019 | |
| 31 40N | 64 10W | North Altantic subtropical gyre | calanoid | 105 | | 0.1008 | 6 | Ivory et al 2019 | 10 year mean; mean of day (90) and night (120) |
| | | Global | copepod | 40.00 | | 0.0384 | 7 | Fernández de Puelles et al 2019 | |
